# TCR convergence in individuals treated with immune checkpoint inhibition for cancer

**DOI:** 10.1101/665612

**Authors:** Timothy Looney, Denise Topacio-Hall, Geoffrey Lowman, Jeffrey Conroy, Carl Morrison, David Oh, Lawrence Fong, Li Zhang

## Abstract

Tumor antigen-driven selection may expand T cells having T cell receptors (TCRs) of shared antigen specificity but different amino acid or nucleotide sequence in a process known as TCR convergence. Substitution sequencing errors introduced by TCRβ (TCRB) repertoire sequencing may create artifacts resembling TCR convergence. Given the anticipated differences in substitution error rates across different next-generation sequencing platforms, the choice of platform could be consequential. To test this, we performed TCRB sequencing on the same peripheral blood mononuclear cells (PBMC) from individuals with cancer receiving anti-CTLA-4 or anti-PD-1 using an Illumina-based approach and an Ion Torrent-based approach. While both approaches found similar TCR diversity, clonality, and clonal overlap, we found that Illumina-based sequencing resulted in higher TCR convergence than with the Ion Torrent approach. Using the latter approach, we found that clonality and convergence independently predicted response and could be combined to improve the accuracy of a logistic regression classifier. These results demonstrate the importance of the sequencing platform in assessing TCRB convergence.

## Introduction

Checkpoint blockade immunotherapy (CPI) may elicit durable anti-tumor responses in a subset of individuals with cancer. Identifying predictive biomarkers to guide treatment selection remains a primary goal of immune-oncology translational research. Owing to limitations in the quantity and quality of available tumor material, and its use in routine PD-L1 immunohistochemistry testing, there is a pressing need to identify non-invasive biomarkers derived from peripheral blood. Within this context tumor mutation burden (TMB) has drawn attention as a potential predictive biomarker for response to CPI, under the premise that it may serve as a surrogate for total neoantigen load and thus the sensitivity of a tumor to immunotherapy. Originally measured from tumor biopsy material, TMB measurements have now been demonstrated from next generation sequencing of peripheral blood cfDNA (Gandara). Unfortunately, accumulating evidence suggests the predictive value of this biomarker may be limited (Goodman), with recent analyses of TMB in mono- or combination CPI for non-small cell lung cancer (NSCLC) indicating an overall Area under the receiver operator characteristic (ROC) curve (AUC) of .60 and .68 respectively, for predicting durable clinical benefit (Rizvi 2018, Hellman 2018), comparable to the accuracy of PD-L1 IHC, while results of Checkmate 026, a study of first-line Nivolumab for NSCLC, revealed no difference in overall survival in patients stratified by TMB (Carbone). Importantly, TMB is unable to identify immunogenic, CPI sensitive tumors having neoantigens other than those derived from non-synonymous mutations, as has been demonstrated by studies of TMB in polyoma virus-associated Merkel cell carcinoma and renal cell carcinomas (Chan).

Motivated by the shortcomings of existing non-invasive biomarkers, here we evaluated the use of peripheral blood TCRB repertoire sequencing as a source of predictive biomarkers for response to CTLA-4 monotherapy for cancer. Previous TCR sequencing studies have evaluated T cell clonal expansion as a stand-alone predictive biomarker, with mixed results (Roh, Voong). One outstanding question is whether TCR sequencing may be used for in silico identification of tumor antigen specific T cells, given that the frequency of such cells could serve as a direct measurement of tumor immunogenicity. Although there are no known methods to predict the antigen specificity of a TCR from nucleotide sequence, we hypothesized that the central role of chronic antigen stimulation in the emergence of cancer would provide a means to infer the presence of tumor antigen specific T cells, given that sustained antigen-driven selection may give rise to convergent T cell receptors having a shared antigen specificity (i.e. identical amino acid sequence) but different nucleotide sequences. Unlike biomarkers relying of the quantification of tumor genetic alterations, TCR convergence: (1) may detect T cell responses to tumor neoantigens beyond those arising from non-synonymous mutations; (2) avoids probabilistic models for prediction of immunogenicity; (3) is sequencing efficient, typically requiring less than 2M reads per sample; and (4) may be measured from the abundant genetic material within the buffy coat fraction of centrifuged peripheral blood to enable liquid biopsy applications.

Despite these advantages, efforts to evaluate TCR convergence may be hampered by the sensitivity of this feature to substitution sequencing errors, which may create artifacts resembling convergent TCRs. To circumvent this issue, here we leveraged the low substitution error rate of the Ion Torrent platform to evaluate convergence as a predictive biomarker for response to anti-CTLA-4 monotherapy in a set of 22 study subjects with cancer. For context, we compared convergence values obtained using this platform to those for the same samples interrogated with Illumina-based TCRB repertoire sequencing. Finally, we examined whether TCR convergence may be combined with measurements of clonal expansion to improve prediction of immunotherapy response.

## Materials and Methods

### Peripheral blood samples

Eight peripheral blood leukocyte (PBL) samples were obtained from longitudinal blood draws from three anti PD-1 treated melanoma study subjects (donor 1: 3 samples; donor 2: 3 samples; donor 3: 2 samples) at the University of California San Francisco (UCSF). The average time between consecutive blood draws was 4 weeks. Baseline (pre-treatment) PBL were collected from 22 cancer study subjects treated with CTLA-4 monotherapy (Ipilimumab) at Roswell Park Cancer Research Institute or UCSF. Samples were collected within one week of administration of the first CTLA-4 dose. Response was evaluated using RECIST criteria.

### TCR sequencing

For the Ion Torrent-based approach, RNA was extracted from cryopreserved buffy coat straws using the Qiagen RNeasy Midi Kit (Qiagen Cat. No. 75144). Extractions were performed over multiple days at two different sites. Purified RNA samples were quantified using Qubit RNA HS Assay Kit (Thermo Fisher Scientific Cat. No. Q32852). The Agilent 2100 Bioanalyzer and Agilent RNA 6000 Nano Kit were used to quantify and evaluate RNA integrity. 25ng of total RNA was reverse transcribed using SuperScript IV VILO Master Mix (Thermo Fisher Scientific Cat. No. 11756050). For each sample, 25ng cDNA was amplified using the Oncomine TCR Beta-LR Assay (Thermo Fisher Scientific Cat. No. A35386) and protocol as described in the Oncomine TCR Beta Assay User Guide MAN0017438 Revision A.0. Libraries were purified with Agencourt AMPure XP beads (Beckman Coulter Cat. No. A63880), washed with 70% ethanol, and eluted in 50uL Low TE buffer. Resulting library samples were diluted 1:100 and quantified using the Ion Library Quantitation Kit (Thermo Fisher Scientific Cat. No. 4468802), then diluted to 25pM with Low TE buffer. Equal volumes from 8 samples at a time were pooled together for sequencing on one Ion 530 chip, followed by analysis via Ion Reporter version 5.10. TCR sequencing with the Illumina-based platform was performed by Adaptive Biotech (Klinger).

### Calculation of TCR convergence and clonality

TCR convergence was calculated as the aggregate frequency of clones sharing a variable gene (excluding allele information) and CDR3AA sequence with at least one other identified clone. For the Oncomine TCRB-LR assay, TCR convergence is pre-calculated from the set of identified clones and provided as a standard output. TCR convergence values in Illumina-based sequencing data were calculated with a custom R script in an identical manner to that for the TCRB-LR assay. Shannon diversity was calculated using the set of clone frequencies (*p*) as indicated below:

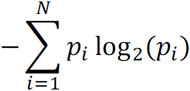

while the normalized Shannon entropy (i.e. evenness) was calculated by dividing the Shannon diversity by log_2_(*N*), where *N* is the total number of detected clones. Clonality was defined as 1 – normalized Shannon entropy.

### Comparison of repertoire features across Illumina and Ion Torrent Datasets

The proportion of overlapping clones between two timepoints within the same subject was evaluated by Jaccard index. Pearson’s product moment correlation coefficient (‘cor.test’ in R ‘stats’ library) was used to compare each repertoire feature across Illumina and Ion Torrent Datasets. Statistical significance was declared based on p<0.05. No multiple testing adjustment was carried out.

### Logistic regression-based prediction of response and model scoring

A logistic regression model with the clinical response status as the binary outcome and TCR convergence and clonality as the predictors was trained using the function ‘train’ from the R ‘caret’ library with the following parameters: method=”glm” and family=”binomial”. Model scores represent the response probability values obtained by applying the caret function ‘predict’, with the trained model and the sample clonality and convergence values as inputs. Model performance was evaluated using the function ‘roc’ from the R ‘pROC’ library. Area under the receiver operator characteristic (ROC) curve (AUC) was calculated using the function ‘auc’ from the same R library. The same functions were used to evaluate the performance of TCR convergence and clonality as stand-alone biomarkers. Cross-validation was performed by training the logistic regression classifier using the trainControl feature of the caret package with the parameters: method=”lgocv”, number=2000, p=.75, classProbs=True, and savePredictions=”final”. Models were separately trained using both TCR convergence and clonality as the predictors, or either feature alone. The optimal score threshold for distinguishing responders from non-responders, and the sensitivity, specificity, and positive predictive value at the optimal threshold, were obtained using the “coords” function from the pROC library with best.method=”youdens” and ret= c(“threshold”, “specificity”, “sensitivity”, “ppv”).

## Results

### Cross-platform analysis of repertoire features

We assessed 8 samples from anti-PD1 treated melanoma subjects using a commercial Illumina-based TCRB sequencing (Adaptive Biotechnologies) and the Ion Torrent based Oncomine TCRB-LR assay. For each sequenced library, we evaluated the number of clones detected, Shannon diversity index, clonality (i.e. 1-normalized Shannon entropy) and the frequency of convergent TCRs (methods). Within each subject from the same sequencing platform, clonal overlap between the samples from any two time points for the same subject was calculated by Jaccard index. We found measurements of Shannon diversity index, clonality, and clonal overlap to be significantly correlated between the two platforms (Figure 1A-C, Pearson’s correlation > .9; p < .001).

**Figure 1.**
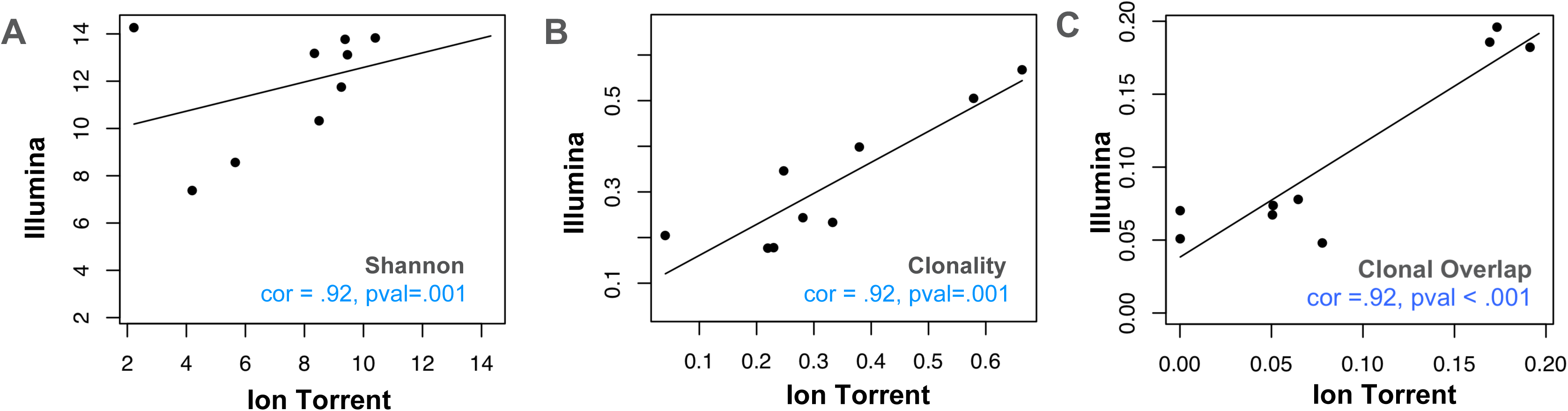
Comparative analysis of repertoire features in samples analyzed via Ion Torrent and Illumina-based assays. Eight peripheral blood leukocyte (PBL) samples derived from three donors were analyzed using the Oncomine TCRB-LR assay (Ion Torrent, X-axis) or Adaptive Biotechnologies TCRB assay (Illumina, Y-axis). Pearson’s correlation coefficient was used to measure the consistency of two platforms with respect to (A) clone diversity (Shannon entropy), (B) clonality (normalized Shannon entropy), and (C) clonal overlap. Blue dashes indicate position of identity line.

### Assessment of TCR convergence

Antigen-driven responses should result in the expansion of multiple T cell clones that recognize this antigen. TCR convergence is calculated as the aggregate frequency of clones sharing a variable gene (excluding allele information) and CDR3AA sequence with at least one other identified clone. An example of a convergent TCR group identified in an individual with melanoma is presented in Figure 2A. For the Oncomine TCRB-LR assay, TCR convergence is pre-calculated from the set of identified clones and provided as a standard output. We hypothesized that the choice of sequencing platform would be consequential for the measurement of TCR convergence given that substitution sequencing errors may mimic TCR repertoire diversity deriving from N-additions and exonucleotide chewback within the V-D and D-J junctions of the CDR3. We found that convergence measurements were not significantly correlated (Figure 2B, Pearson’s correlation = .57; p = .14), with 8 of 8 Illumina libraries showing higher TCR convergence compared to the corresponding Ion Torrent libraries (p = 0.002, Wilcoxon signed rank test).

**Figure 2.**
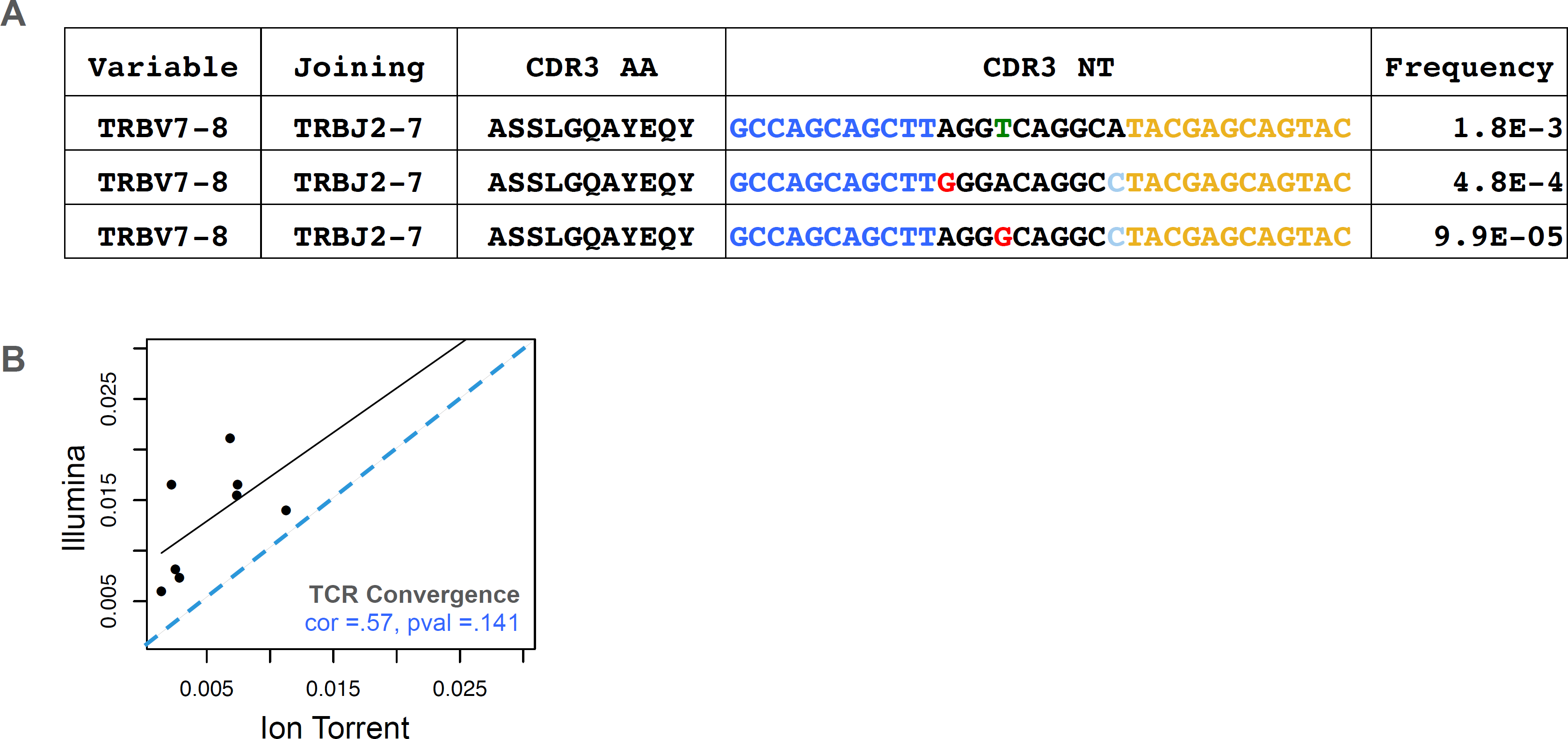
Assessment of TCR convergence. (A) Example of a convergent TCR group detected in the peripheral blood of an individual with melanoma. This group consists of three TCRβ clones that are identical in TCRβ amino acid space but have distinct CDR3 NT junctions owing to differences in non-templated bases at the V-D-J junction. Blue indicates bases contributed by the variable gene while yellow indicates bases contributed by the joining gene. Red arrows indicate positions where clones differ. Substitution sequencing errors and PCR errors can create artifacts that resemble convergent TCRs. (B) TCR convergence, calculated as the aggregate frequency of clones sharing an amino acid sequence with at least one other clone. Blue dashes indicate position of identity line.

**Figure 3.**
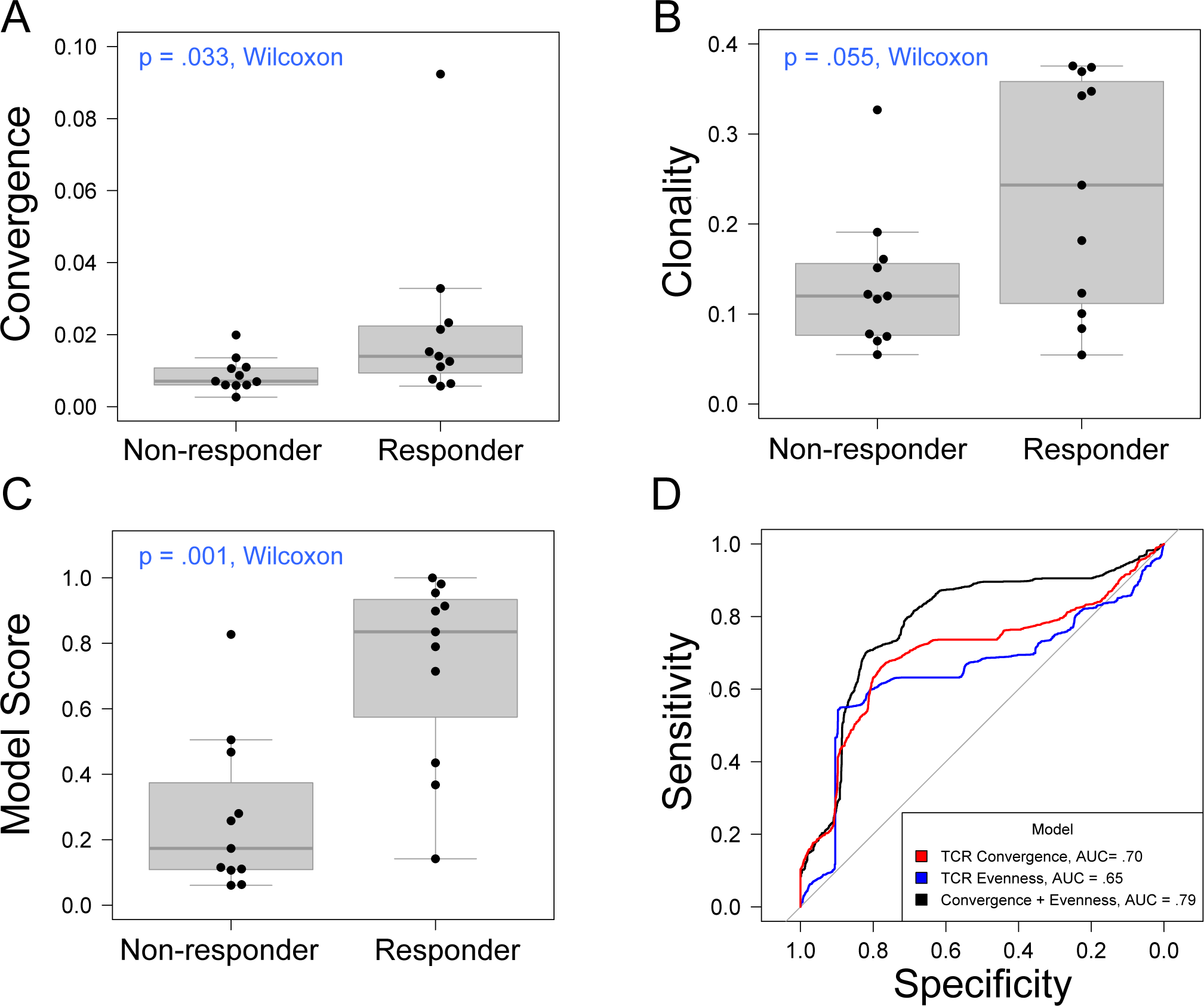
Association between clinical outcomes and TCR convergence. (A) TCR convergence and (B) clonality for responders (N=11) and non-responders (N=11) to CTLA-4 blockade for cancer. TCR clonality is calculated as 1 - the normalized Shannon entropy of clone frequencies. Convergent TCR frequency was calculated as described in methods. All cancer types were included in the analysis. (C) Response probability scores from a logistic regression classifier trained using TCR clonality and convergence as features to predict response to immunotherapy. Score indicates likelihood that a sample is a responder. (D) Receiver operator characteristic curves derived from leave-group-out cross validation analysis of models using clonality, convergence, or the combination of clonality and convergence to predict immunotherapy response. ROC curves represent the average model performance following 2000 random train-test splits, where 75% of the dataset was used to train the model followed by testing on the remaining 25%. The combination of TCR clonality and convergence shows better performance (AUC = .89) than models using TCR convergence and clonality alone (AUC of .70 and .65, respectively).

### Association with response to CTLA-4 monotherapy

We next used the Oncomine TCRB-LR assay to evaluate TCR convergence as a predictive biomarker for response to CTLA-4 blockade in a cohort of 22 individuals with RECIST graded response annotations (11 responders, 11 non-responders) representing three major cancer types: clear cell adenocarcinoma (2 responders, 0 non-responder), melanoma (7 responders, 6 non-responders) and prostate cancer (2 responders, 4 non-responders). Response was defined as stable disease, partial response, or complete response following immunotherapy. Cancer annotations and repertoire features for this cohort are presented in Table 1. We found TCR convergence to be elevated in those who had an objective response to immunotherapy (p = .033, Wilcoxon sum rank test, Figure 2A) and could discriminate responders from non-responders with an AUC of .77. Given previous reports on the potential biomarker value of T cell clonal expansion, we next asked whether clone clonality values differed between responders and non-responders. We observed a trend towards higher clonality in those who responded to immunotherapy (p= .055, Wilcoxon sum rank test, Figure 2B; AUC= .74). Given that clonal expansion and TCR convergence measure independent repertoire features, we trained a logistic regression classifier using TCR convergence and clonality as the two model features to test whether they might be combined to improve the prediction of response (methods). We found that the combination of convergence and clonality improved the prediction of response (Wilcoxon sum rank test p = .001, model response probability score for responders vs. non-responders; Figure 2C, AUC= .89). Finally, to evaluate model robustness, we performed repeated leave-group-out cross validation of the two feature models, and compared performance to the models using TCR convergence or clonality as a sole predictor of response. We found the two-feature model to outperform those using a single feature, achieving a specificity of .82, sensitivity of .71, and positive predictive value of .80 at the optimal threshold score, as determined by Youden’s J (Figure 2D and methods).

**Table 1.** Cancer type and summary repertoire features for 22 individuals receiving CTLA-4 monotherapy. Summary repertoire features and sample annotations for cohort. Each individual was profiled via the Oncomine TCRB-LR Assay at a single baseline timepoint using 25ng of cDNA derived from PBL total RNA. Repertoire feature values indicate the average and range for responders and non-responders.

| Category | Subdefinition | Responder | Non-responder |
| --- | --- | --- | --- |
| Cancer Type | Prostate | 2 | 4 |
|  | Melanoma | 7 | 6 |
|  | Adenocarcinoma | 2 | 0 |
|  | Not Indicated | 0 | 1 |
|  | Total | 11 | 11 |
| Repertoire Features | Clones Detected | 32916 (5168-56231) | 30015 (5894-58222) |
|  | TCR Convergence | 0.022 (.006- .092) | .008 (.002-.019) |
|  | Clonality | .24 (.055 - .376) | .133 (.055-.327) |

## Discussion

The majority of TCRB sequencing data published to date has been generated using the Illumina platform, which has inherent substitution sequencing errors (Schirmer). Platforms having a lower substitution error rate could produce more suitable data. We hypothesized that the low substitution error rate of Ion Torrent platform might be critical for the measurement of convergence (Looney, Merriman). Indeed, in a side-by-side comparison of samples analyzed using Illumina and Ion Torrent based sequencing, we found measurements of TCR clonality, diversity and clonal overlap to be consistent across platforms, while TCR convergence values were not significantly correlated. In reviewing literature, we note other reports of TCR convergence in Illumina based data that are significantly higher than those we observe in Ion Torrent data (Ruggiero). Taken together, these results suggest the choice of sequencing platform may be consequential for biomarker applications of TCR convergence, and raise the possibility that TCR convergence may have been overlooked as a predictive biomarker owing to obfuscating platform specific noise.

Here we report elevated TCR convergence in baseline peripheral blood of those who respond to CTLA-4 blockade for cancer. We define TCR convergence as the aggregate frequency of T cell clones within an individual that share a variable gene and CDR3AA sequence with at least one other clone, but differ at the nucleotide level. This definition can be contrasted with instances in literature where this term has been used to refer to TCRB or IGH amino acid sequences found in more than one individual (Venturi 2006, Venturi 2008b, Parmeswarwan) (i.e. “public” rearrangements) or instances where researchers attempt to identify functionally equivalent TCRs that differ in amino acid space (Dash). Compared to the latter approach, we have adopted a stringent definition of convergence, with the goal of minimizing the false positive rate for the detection of convergent TCRs. Our approach builds upon the notion that false positive convergent events, either owing to the grouping of functionally dissimilar clones, or the presence of artifactual clones deriving from residual substitution errors, have the potential to conceal meaningful signal in TCR repertoire data, a possibility exacerbated by the rarity of bone fide convergence events. Consequently, we note that the frequency of convergent TCRs reported here may underestimate the frequency of functionally equivalent T cell clones.

One proposed advantage of TCR convergence as a biomarker is its ability to detect T cell responses to tumor neoantigens beyond those arising from non-synonymous mutations. Although this dataset is small, we find that convergence values could discriminate responders from non-responders with significant accuracy (AUC = .77), comparing favorably to the historical performance of tumor mutation burden as a biomarker. The extent to which elevated TCR convergence is a feature of CPI sensitive tumors of other cancer types will be clarified by ongoing studies involving larger cohorts.

We hypothesize that T cells having convergent TCRs are likely to target tumor-associated antigens in those with cancer. However, these data do not shed light on the phenotype of such tumor antigen specific T cells. Given that chronic antigen stimulation may give rise to exhausted or dysregulated T cells having distinct expression levels of cell surface receptors, including inhibitory receptors, (Kurachi), it is possible that FACS-based methods may be used to enrich for T cells having convergent TCRs. This possibility may be relevant given recent reporting of a phenotypically abnormal PD-1 high intratumoral T cell population in NSCLC patients treated with PD-1 blockade, the frequency of which was found to be predictive of immunotherapy response. (Thommen).

One question arising from this work is whether additional insight would be gained by analysis of the unseen TCRα (TCRA) chains of T cells having convergent TCRB chains. Accepting that members of a convergent TCR group share antigen specificity, one can infer that either: 1) the unseen TCRA chains are functionally identical and help determine the antigen specificity of the receptor; or 2) convergent TCRs are TCRB chain dominant, while the TCRA chains play an accessory or stabilizing role but do not affect the antigen specificity of the receptor. Given that the unsequenced TCRA chains of group members are likely to differ in sequence space owing to the random nature of VDJ recombination, the latter case may be more likely. In this scenario, the full length TCRB chains of convergent TCRs could be paired with generic TCRA chains to efficiently generate tumor antigen specific TCRs.

Finally, although it is not a focus of this study, the link between chronic antigen stimulation and autoimmunity suggests that TCR convergence may also have application to the identification of mechanistic or predictive biomarkers for autoimmune disease. To this end we note that TCR convergence has been identified as a repertoire feature of salivary gland inflammatory lesions in individuals with Sjogren’s syndrome (Joachims).

## References

1. Gandara D, Paul S, Kowanetz M, Schleifman E, Zou W, Li Y et al. Blood based tumor mutational burden as a predictor of clinical benefit in non-small-cell lung cancer patients treated with atezolizumab. Nature Medicine 2018; 24: 1441–1448.

2. Goodman AM, Kato S, Bazhenova L, Patel SP, Frampton GM, Miller V et al. Tumor mutational burden as an independent predictor of response to immunotherapy in diverse cancers. Mol Cancer Ther 2017; 11:2598–2608.

3. Rizvi H, Sanchez-Vega F, La K, Chatila W, Jonsson P, Halpenny D et al. Molecular determinants of response to anti-programmed cell Death (PD)-1 and Anti-Programmed Death-Ligand 1 (PD-L1) Blockade in Patients With Non-Small-Cell Lung Cancer Profiled With Targeted Next-Generation Sequencing. J Clin Oncol 2018; 36:633–641.

4. Hellmann MD, Nathanson T, Rizvi H, Creelan BC, Sanchez-Vega F, Ahuja A et al. Genomic features of response to combination immunotherapy in patients with advanced small-cell lung cancer. Cancer Cell 2018; 33:843–852.

5. Carbone D, Reck M, Pas-Ares L, Creelan B, Horn L, Steins M et al. First-line Nivolumab in stage IV or recurrent non-small cell lung cancer. N Engl J Med 2017; 376: 2415–2426.

6. Chan TA, Tarchoan M, Jaffee E, Swanton C, Quezada SA, Steinzinger A et al. Development of tumor mutation burden as an immunotherapy biomarker: utility for the oncology clinic. Ann Oncol 2018; doi:10.1093

7. Roh W, Pei-Ling C, Reuben A, Spencer C, Prieto P, Miller J et al. Intergrated molecular analysis of tumor biopsies on sequential CTLA-4 and PD-2 blockade reveals markers of response and resistance. Sci Transl Med 2017; 9: doi:10.1126.

8. Voong K, Feliciano J, Becker D and Levy B. Beyond PD_L1 testing-emerging biomarkers for immunotherapy in non-small cell lung cancer. Ann Transl Med 2017; 5:376.

9. Klinger M, Moorhead M, Weng L, Zheng J, Faham M, Combining next-generation sequencing and immune assays: A novel method for identification of antigen-specific T cells. PLOS One, 2013. 8(e74231).

10. Schirmer M, D’Amore R, Ijaz UZ, Hall N and Quince C. Illumina error profiles: resolving fine-scale variation in metagenomic sequencing data. BMC Bioinformatics 2016; 17:125.

11. Looney T, Duose D, Lowman G, Linch E, Hajjar J, Topacio-Hall D et al. Haplotype analysis of the TRB locus by TCRB repertoire sequencing. bioRxiv 2018; doi:10.1101/406157.

12. Merriman B, Ion Torrent R&D Team, and Rothberg JM. Progress in Ion Torrent semiconductor chip based sequencing. Electrophoresis 2010; 33: 3397–3417.

13. Venturi V, Kedzierska K, Price DA, Doherty PC, Douek DC, Turner SJ et al. Sharing of T cell receptors in antigen-specific responses is driven by convergent recombination. Proc Natl Acad Sci USA 2006; 103:18691–6.

14. Venturi V, Chin HY, Asher TE, Ladell K, Scheinberg P, Borstein E et al. TCR beta-chain sharing in human CD8+ T cell responses to cytomegalovirus and EBV. J Immunol 2008; 181:7853–62.

15. Parameswaran P, Liu Y, Roskin KM, Jackson KK, Dixit VP, Lee JY et al. Convergent antibody signatures in human dengue. Cell Host Microbe 2013; 13:691–700.

16. Dash P, Fiore-Garland AJ, Hertz T, Wang GC, Gharma S, Souquette A et al. Quantifiable predictive features define epitope-specific T cell receptor repertoires. Nature 2017; 547:89–93.

17. Wherry J and Kurachi M. Molecular and cellular insights into T cell exhaustion. Nat Rev Immunol 2015; 15:486–499.

18. Thommen D, Koelzer V, Herzig P, Roller A, Trefny M, Dimeloe S et al. A transcriptionally and functionally distinct PD-1+ CD8+ T cell pool with predictive potential in non-small-cell lung cancer treated with PD-1 blockade. Nature Medicine 2018; 24: 994–1004.

19. Ruggiero E, Nicolay JP, Fronza R, Arens A, Paruzynski A, Nowrouzi A et al. High-resolution analysis of the human T-cell receptor repertoire. Nat Commun 2015; 6:8081.

20. Joachims M, Leehan KM, Lawrence C, Pelikan RC, Moore JS, Pan Z et al. Single-cell analysis of glandular T cell receptors in Sjogren’s syndrome. JCI Insight 2016; 3: pii: e85609.

